# From fitting the average to fitting the individual: A cautionary tale for mathematical modelers

**DOI:** 10.1101/2021.08.03.454882

**Authors:** Michael C. Luo, Elpiniki Nikolopoulou, Jana L. Gevertz

## Abstract

An outstanding challenge in the clinical care of cancer is moving from a one-size-fits-all approach that relies on population-level statistics towards personalized therapeutic design. Mathematical modeling is a powerful tool in treatment personalization, as it allows for the incorporation of patient-specific data so that treatment can be tailor-designed to the individual. Herein, we work with a mathematical model of murine cancer immunotherapy that has been previously-validated against the average of an experimental dataset. We ask the question: what happens if we try to use this same model to perform personalized fits, and therefore make individualized treatment recommendations? Typically, this would be done by choosing a single fitting methodology, and a single cost function, identifying the individualized best-fit parameters, and extrapolating from there to make personalized treatment recommendations. Our analyses show the potentially problematic nature of this approach, as predicted personalized treatment response proved to be sensitive to the fitting methodology utilized. We also demonstrate how a small amount of the right additional experimental measurements could go a long way to improve consistency in personalized fits. Finally, we show how quantifying the robustness of the average response could also help improve confidence in personalized treatment recommendations.

**Author summary:** As we enter the era of healthcare where personalized medicine becomes a more common approach to treating cancer patients, harnessing the power of mathematical models will only become more essential. Using a preclinical dataset on cancer immunotherapy, we explore the challenges and limitations that arise when trying to move from fitting and making predictions for the population-level average, to fitting and making predictions for an individual. We find that the standard of approach of picking a single fitting methodology and a single cost function may end up having limited predictive value when applied to individual data. We also show how having a small amount of the right additional experimental data, and establishing the robustness of average treatment response, can help improve confidence in personalized model predictions.

## Introduction

The conventional approach for developing a cancer treatment protocol relies on measuring average efficacy and toxicity from population-level statistics in randomized clinical trials [1–3]. However, it is increasingly recognized that heterogeneity, both between patients and within a patient, is a defining feature of cancer [4, 5]. This inevitably results in a portion of cancer patients being over-treated and suffering toxicity consequences from the standard-of-care dose, and another portion being under-treated and not benefiting from the expected efficacy of the treatment [6].

For these reasons, in the last decade there has been much interest in moving away from this ‘one-size-fits-all’ approach to cancer treatment and towards personalized therapeutic design (also called predictive or precision medicine) [1, 2, 7]. Collecting patient-specific data has the potential to improve treatment response to chemotherapy [6, 8–11], radiotherapy [12–14], and targeted molecular therapy [11, 15–17]. However, it has been proposed that personalization may hold the most promise when it comes to immunotherapy [18]. Immunotherapy is an umbrella term for methods that increase the potency of the immune response against cancer. Unlike other treatment modalities that directly attack the tumor, immunotherapy depends on the interplay between two complex systems (the tumor and the immune system), and therefore may exhibit more variability across individuals [18].

Mathematical modeling has become a valuable tool for understanding tumor-drug interactions. However, just as clinical care is guided by standardized recommendations, most mathematical models are validated based on population-level statistics from preclinical or clinical studies [19]. To truly realize the potential of mathematical models in the clinic, these models must be individually parameterized using measurable, patient-specific data. Only then can modeling be harnessed to answer some of the most pressing questions in precision medicine, including selecting the right drug for the right patient, identifying optimal drug combinations for a patient, and prescribing a treatment schedule that maximizes efficacy while minimizing toxicity.

Efforts to personalize mathematical models have been undertaken to understand glioblastoma treatment response [20, 21], to identify optimal chemotherapeutic and granulocyte colony-stimulating factor combined schedules in metastatic breast cancer [22], to identify optimal maintenance therapy chemotherapeutic dosing for childhood acute lymphoblastic leukemia [9], and to identify optimized doses and dosing schedules of the chemotherapeutic everolimus with the targeted agent sorafenib for solid tumors [23]. Interesting work has also been done in the realm of radiotherapy, where individualized head and neck cancer evolution has been modeled through a dynamic carrying capacity informed by patient response to their last radiation dose [24].

Beyond these examples, most model personalization efforts have focused on prostate cancer, as prostate-specific antigen is a clinically measurable marker of prostate cancer burden [25] that can be used in the parameterization of personalized mathematical models. The work of Hirata and colleagues has focused on the personalization of intermittent androgen suppression therapy using retrospective clinical trial data [26, 27]. Other interesting work using clinical trial data has been done by Agur and colleagues, focusing on individualizing a prostate cancer vaccine using retrospective phase 2 clinical trial data [25, 28], as well as androgen deprivation therapy using data from an advanced stage prostate cancer registry [29]. Especially exciting work on personalizing prostate cancer has been undertaken by Gatenby and colleagues, who used a mathematical model to discover patient-specific adaptive protocols for the administration of the chemotherapeutic agent abiraterone acetate [30]. Among the 11 patients in a pilot clinical trial treated with the personalized adaptive therapy, they observed the median time to progression increased to at least 27 months as compared to 16.5 months observed with standard dosing, while also using a cumulative drug amount that was 47% less than the standard dosing [17].

Despite these examples, classically mathematical models are not personalized, but are validated against the average of experimental data. In particular, modelers choose a single fitting methodology, a single cost function to minimize, and find the best-fit parameters to the average of the data. Using the best-fit parameters and the mathematical model, treatment optimization can be performed. Recognizing the limitations of this approach in describing variable treatment response across populations, modelers have begun employing virtual population cohorts [31, 32]. There is much value in this population-level approach to study variability, but it is not equivalent to looking at individualized treatment response.

In this work, we explore the consequences of performing individualized fits using a minimal mathematical model previously-validated against the average of an experimental dataset. In Materials and methods, we describe the preclinical data collected by Huang et al. [33] on a mouse model of melanoma treated with two forms of immunotherapy, and our previously-developed mathematical model that has been validated against population-level data from this trial [34]. Individual mouse volumetric time-course data is fit to our dynamical systems model using two different approaches detailed in Materials and methods: the first fits each mouse independent of the other mice in the population, whereas the second constrains the fits to each mouse using population-level statistics. In Results and Discussion, we demonstrate that the treatment response identified for an individual mouse is *sensitive to the fitting methodology utilized*. We explore the causes of these predictive discrepancies and how robustness of the optimal-for-the-average treatment protocol influences these discrepancies. We conclude with actionable suggestions for how to increase our confidence in mathematical predictions made from personalized fits.

## Materials and methods

### Data Set

The data in this study considers the impact of two immunotherapeutic protocols on a murine model of melanoma [33]. The first protocol uses oncolytic viruses (OVs) that are genetically engineered to lyse and kill cancer cells. In [33] the OVs are immuno-enhanced by inserting transgenes that cause the virus to release 4-1BB ligand (4-1BBL) and interleukin (IL)-12, both of which result in the stimulation of the tumor-targeting T cell population [33]. The preclinical work of Huang et al. has shown that oncolytic viruses carrying 4-1BBL and IL-12 (which we will call Ad/4-1BBL/IL-12) can cause tumor debulking via virus-induced tumor cell lysis, and immune system stimulation from the local release of the immunostimulants [33].

The second protocol utilized by Huang et al. are dendritic cell (DC) injections. DCs are antigen-presenting cells that, when exposed to tumor antigens ex vivo and intratumorally injected, can stimulate a strong adaptive immune response against cancer cells [33]. Huang et al. showed that combination of Ad/4-1BBL/IL-12 with DC injections results in a stronger antitumor response than either treatment individually [33]. Volumetric trajectories of individual mice treated with Ad/4-1BBL/IL-12, along with the average trajectory, is shown in Fig. 1.

**Fig 1.**
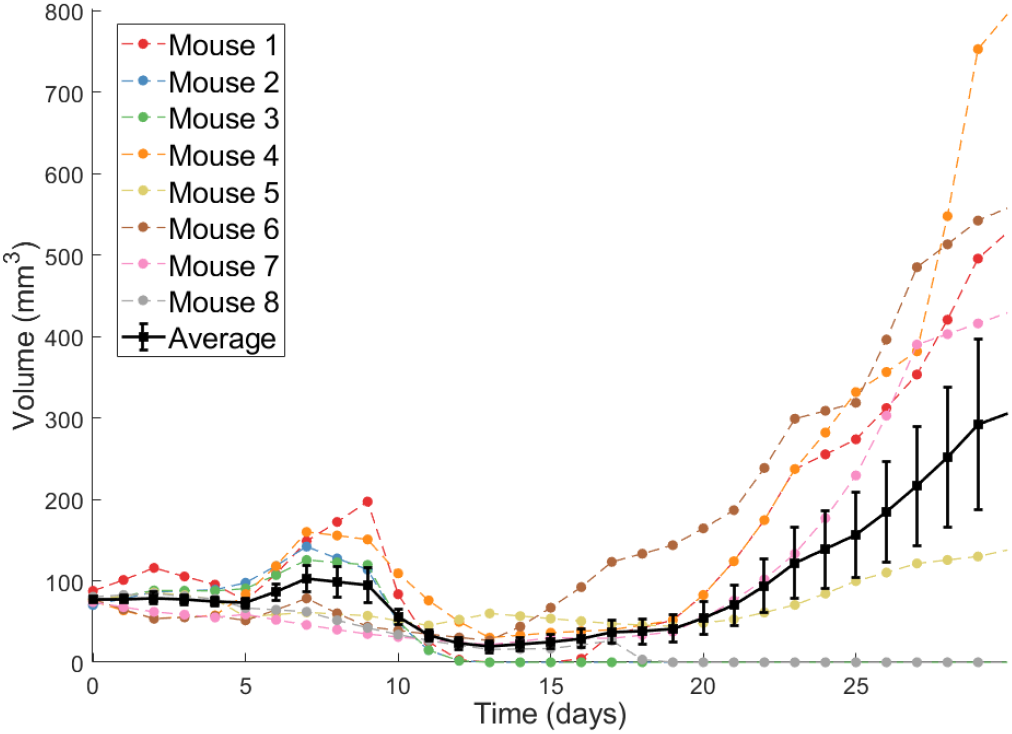
Individual volumetric trajectories are shown for eight mice treated with Ad/4-1BBL/IL-12. The average, with standard error bars, is also shown in black [33].

### Mathematical Model

Our model contains the following five ordinary differential equations:

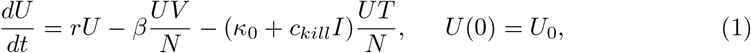

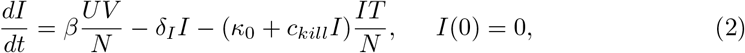

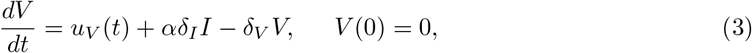

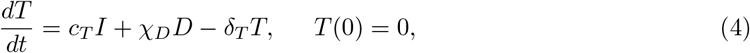

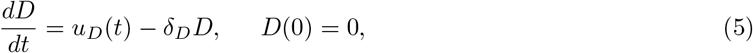

where *U* is the volume of uninfected tumor cells, *I* is the volume of OV-infected tumor cells, *V* is the volume of free OVs, *T* is the volume of tumor-targeting T cells, *D* is the volume of injected dendritic cells, and *N* is the total volume of cells (tumor cells and T cells) at the tumor site. When all parameters and time-varying terms are positive, this models captures the effects of tumor growth and response to treatment with Ad/4-1BBL/IL-12 and DCs [34]. By allowing various parameters and time-varying terms to be identically zero, other treatment protocols tested in Huang et al. [33] can also be described.

This model was built in a hierarchical fashion, details of which have been described extensively elsewhere [31, 34–36]. Here, we briefly summarize the full model. Uninfected tumor cells grow exponentially at a rate *r*, and upon being infected by an OV convert to infected cancer cells at a density-dependent rate *βUV/N*. These uninfected cells get lysed by the virus or other mechanisms at a rate of *δ*_*I*_, thus acting as a source term for the virus by releasing an average of *α* free virions into the tissue space. Viruses decay at a rate of *δ*_*V*_.

The activation/recruitment of tumor-targeting T cells can happen in two ways: 1) stimulation of cytotoxic T cells due to 4-1BBL or IL-12 (modeled through *I*, at a rate of *c*_*T*_, as infected cells are the ones to release 4-1BBL and IL-12), and 2) production/recruitment due to the externally-primed dendritic cells at a rate of *χ*_*D*_. These tumor-targeting T cells indiscriminately kill uninfected and infected tumor cells, with the rate of killing that depends on IL-12 and 4-1BBL production (again, modeled through *I* in the term (*κ*_0_ + *c*_*kill*_*I*)), and they can also experience natural death at a rate of *δ*_*T*_. The time-dependent terms, *u*_*V*_ (*t*) and *u*_*D*_(*t*), represent the source of the drug and are determined by the delivery and dosing schedule of interest.

### Fitting Methodologies

#### Independently Fitting Individuals

Our first attempt at individualized fitting is to find the parameter set that minimizes the *L*^2^-norm between the model and the individual mouse data:

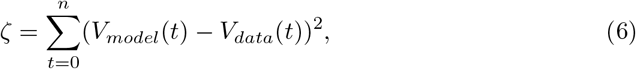

where *V*_*model*_(*t*) = *U* (*t*) + *I*(*t*) is the volumetric output predicted by our model in eqns. (1)-(5), *V*_*data*_(*t*) represents the volumetric data for an individual mouse, and *n* is the number of days for which tumor volume is measured in the experiments.

To independently fit an individual mouse, parameter space is first quasi-randomly sampled using high-dimensional Sobol’ Low Discrepancy Sequences (LDS). LDS are designed to give rise to quasi-random numbers that sample points in space as uniformly as possible, while also (typically) having faster convergence rates than standard Monte Carlo sampling methods [37]. After the best-fit parameter set has been selected among the 10^6^ randomly sampled sets chosen by LDS, the optimal is refined using simulated annealing [38]. Having observed that the landscape of the objective function near the optimal parameter set does not contain local minima, we randomly perturb the LDS-chosen parameter set, and accept any parameter changes that decrease the value of the objective function - making the method equivalent to gradient descent. This random perturbation process is repeated until no significant change in *ζ* can be achieved, which we defined as the relative change in *ζ* for the last five accepted parameter sets being less than 10^−6^. We call this final parameter set the optimal parameter set.

It is important to note that, by approaching fitting in this way, the parameters for Mouse *i* depend only the volumetric data for Mouse *i*; that is, the volumetric data for the other mice are not accounted for.

#### Fitting Individuals with Population-Level Constraints

Nonlinear mixed effects (NLME) models incorporate fixed and random effects to generate models to analyze data that are non-independent, multilevel/hierarchical, longitudinal, or correlated [39]. Fixed effects refer to parameters that can generalize across an entire population. Random effects refer to parameters that differ between individuals that are randomly sampled from a population.

The mixed effects model we will utilize is of the form:

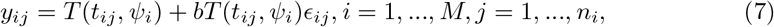

where *y*_*ij*_ is the predicted tumor volume at each day *j* for each individual *i* (that is, at time *t*_*ij*_), *M* = 8 is the number of mice, *n*_*i*_ = 31 is the number of observations per mouse, *ψ*_*i*_ is the parameter vector for the structural model for each individual, and *ϵ*_*ij*_ is a variable describing random noise. Here we made the assumption that the error is a scalar value proportional to our structural model.

Typically, NLME models attempt to maximize the likelihood of the parameter set given the available data. There does not exist a general closed-form solution to this maximization problem [40], so numerical optimization is often needed to find a maximum likelihood estimate. In this work, we employ Monolix [41], which uses a Markov Chain Monte Carlo method to find values of the model parameters that optimize the likelihood function. To implement NLME in Monolix, we first processed and arranged our experimental data consisting of tumor volume and dosing schedule in a Monolix-specified spreadsheet. To avoid predictive errors, we censored the data, as detailed in [41]. More specifically, all tumor volumes less than 1 mm^3^ were set to 0. This was done to prevent over-fitting to these data points at the expense of the rest of the data.

We assume that each parameter *ψ*_*i,k*_ ∈ *ψ*_*i*_ is lognormally distributed with mean 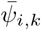 and standard deviation *ω*_*i,k*_:

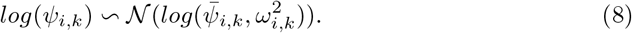

Based on previous fits to the average of the data in [36], we used the following set of initial guesses for the population parameters:

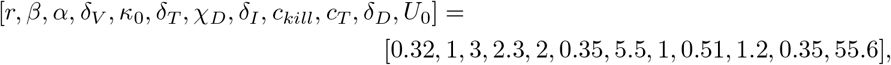

with the initial standard deviations chosen as:

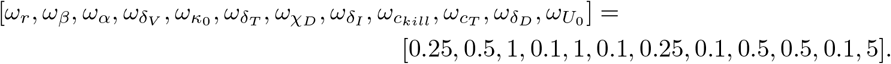

### Practical Identifiability via the Profile Likelihood Method

It is well-established that estimating a unique parameter set for a mathematical model can be challenging due to the limited availability of often noisy experimental data [42]. A non-identifiable model is one in which multiple parameter sets give “good” fits to the experimental data. Here, we will study the practical identifiability of our system in eqns.(1) - (5) using the profile likelihood approach [43, 44].

A single parameter is profiled by fixing it across a range of values, and subsequently fitting all other model parameters to the data [42]. To execute the profile likelihood method, let *p* be the vector that contains all parameters of the model, *θ* be one parameter of interest contained in the vector *p*. The profile likelihood *PL* for the parameter *θ* is defined in [45] as:

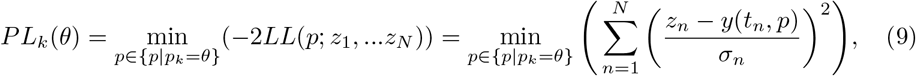

where *z*_*n*_ for *n* = 1, …, *N* is the measured data that is assumed to follow a normal distribution with mean *y*(*t*_*n*_, *p*) and variance *σ*^2^, and *LL*(*p*; *z*_1_, …*z*_*N*_) is the log of the likelihood function. The likelihood function represents the likeliness of the measured data *z*_*n*_ given a model with parameters *p* [46]. The profile likelihood curve for any parameter of interest *θ* is found using the following process:

1. Determine a range for the parameter values of *θ*.
2. Fix *θ* = *θ*^∗^ at a value in the range.
3. For the fixed value in step 2 we fit the parameters 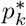 and obtain the best-fit values by minimizing the objective function defined in eqn. (9).
4. Evaluate the objective function at those optimum values for the fixed value of *θ*^∗^.
5. Repeat the process described in steps 2-4 for a discrete set of values in the range of the parameter *θ*. This yields to the profile likelihood function for the parameter *θ*.

Once *PL*(*θ*) is determined, the confidence interval for *θ* at a level of significance *α* can be computed using:

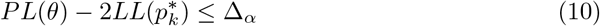

where Δ_*α*_ denotes the *α* quantile of the *χ*^2^ distribution with *df* degrees of freedom (which represents the number of fit model parameters when calculating *PL*(*θ*)) [42]. We use *α* = 0.95 for a 95% confidence interval. The intersection points between the threshold 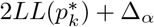 and *PL*(*θ*) result in the bounds of the confidence interval. A parameter is said to be practically identifiable if the shape of the profile likelihood plot is close to quadratic on a finite confidence interval [47]. Otherwise, a parameter is said to be practically unidentifiable.

## Results and Discussion

### Personalized Fits

The individual mouse data in response to treatment with Ad/4-1BBL/IL-12 + DCs [33] is fit using the two methodologies discussed previously: 1) quasi-Monte-Carlo method with simulated annealing in which each mouse is fit independently (which we will call the “QMC” method for short), and 2) nonlinear mixed effects modeling in which population-level statistics constrain individual fits. In Fig. 2, we can see the best-fit for each mouse using the two fitting approaches.

**Fig 2.**
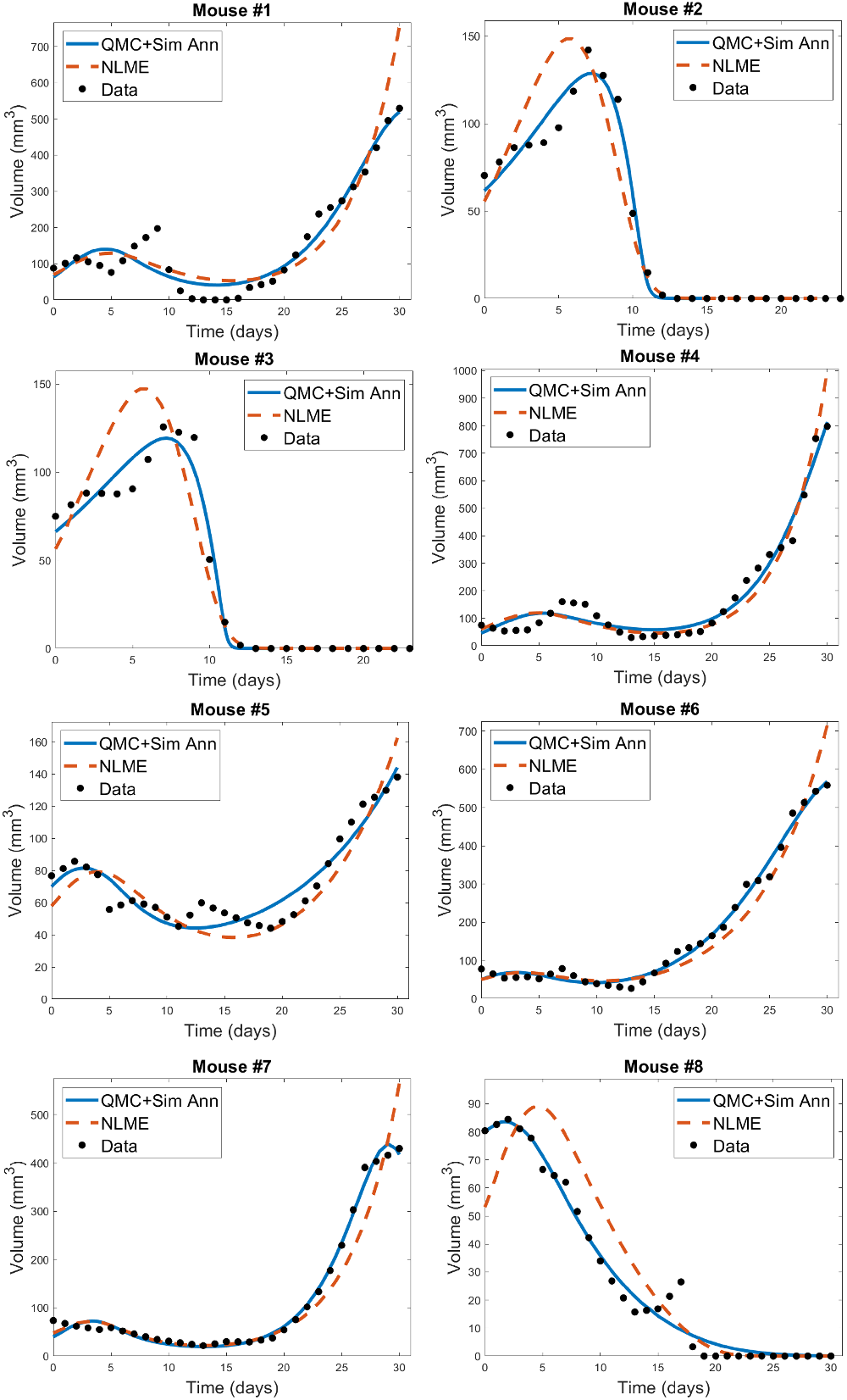
Best-fit for each mouse treated with Ad/41BBL/IL-12 and DCs in the order VDVDVD at a dose of 2.5 × 10^9^ OVs and 10^6^ DCs [33]. The QMC fits (in which each mouse is treated independently of the others) are shown in blue, and the NLME fits are shown in red. The experimental data (black) is also provided on each plot.

We observe that for each mouse, the QMC algorithm results in a fit that more accurately captures the dynamics in the experimental data. The differences between the two fitting methodologies explain why this is occurring. NLME assumes each parameter is sampled from a lognormal distribution whose mean and variance are determined by the full population of mice. The estimated lognormal distributions for each model parameter are shown in Fig. S1. On the other hand, the QMC algorithm fits each mouse independently, and the only constraint imposed on the parameters is a nonnegativity constraint. This allows the QMC algorithm to explore a much larger region of parameter space, resulting in better fits. The downside, as we will show, is that the QMC algorithm may be selecting parameters that are not biologically realistic.

We have established that the parametric constraints across the two fitting methodologies explain the goodness-of-fit differences seen in Fig. 2. However, this does not tell us *which* parameters vary across fitting methodologies and which, if any, are conserved. In Fig. 3 and S2 we show the best-fit parameter value for each mouse and fitting methodology *relative to the best-fit parameter value for the average mouse*. For example, the best-fit value of the tumor growth rate *r* to the average of the control data has been shown to be *r* = 0.3198 [34]. Since Mouse 1 has a relative value of 1.0916 when fitting is done using QMC, the value of *r* predicted for that Mouse is 9.16% larger than the value for the average mouse, meaning QMC predicts *r* = 0.3491 for Mouse 1. On the other hand, the relative value is 0.7512 when fitting is done using NLME, meaning the predicted value is *r* = 0.2402, which is 24.88% less than the value for the average mouse.

**Fig 3.**
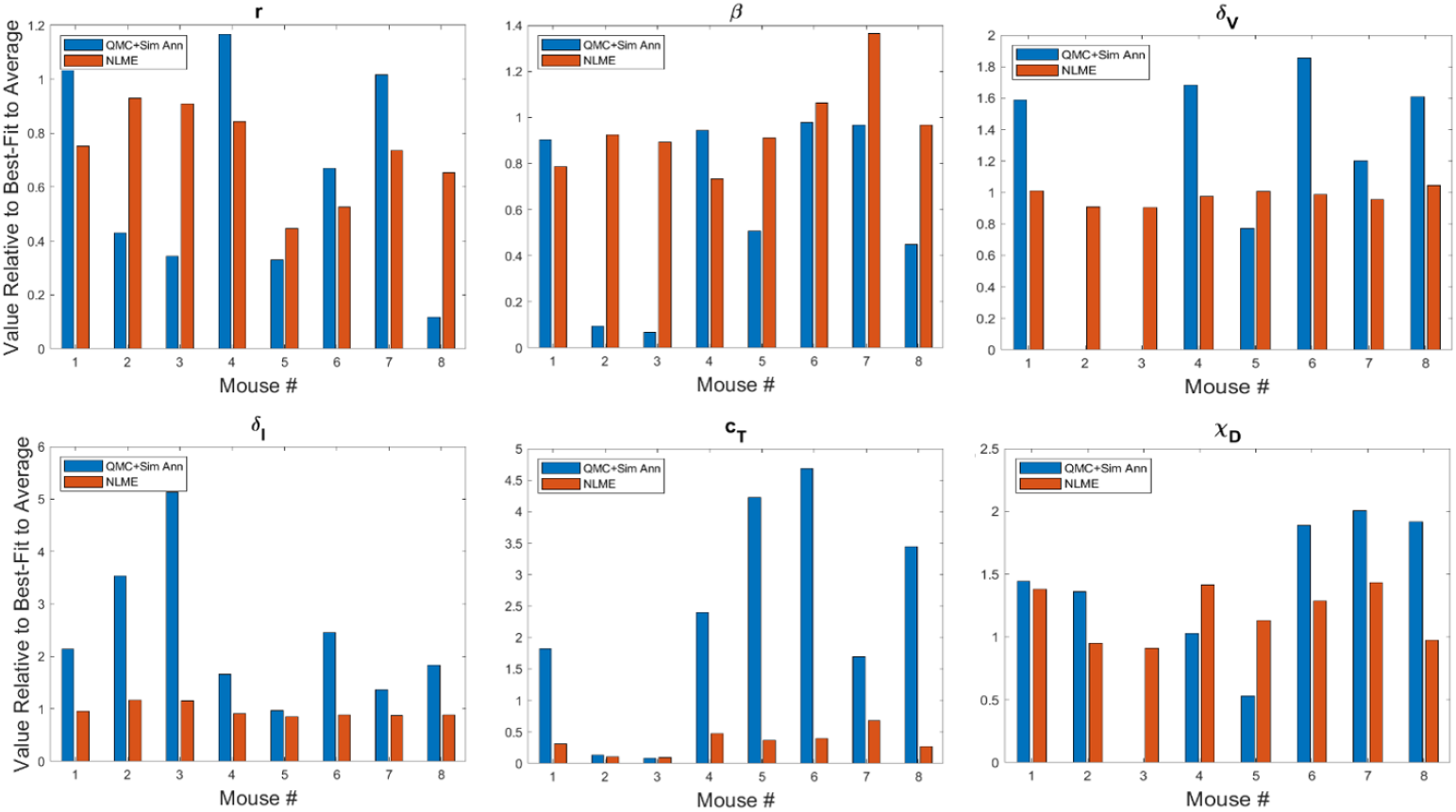
Best-fit values of tumor growth rate parameter *r*, virus infectivity parameter *β*, viral decay rate *δ*_*V*_, infected cell lysis rate *δ*_*I*_, T cell stimulation term by immunostimulants *c*_*T*_, and T cell stimulation term by DCs *χ*_*D*_. The best-fit values are shown for each mouse and are presented relative to the best-fit value of the parameter in the average mouse [34]. Therefore, a value of 1 means the parameter value is equal to that in the average mouse, less than 1 is a smaller value, and greater than 1 is a larger value. Values for other model parameters are shown in Fig. S2.

We observe that while the value a parameter can take on across methodologies is usually of the same order of magnitude, substantial differences can exist across methodologies in essentially all parameters. Generally speaking, NLME-associated parameters exhibit smaller variations from the best-fit parameter for the average mouse. This occurs because the full population of mice constrain the lognormal distribution that each parameter is sampled from.

Compare this to the QMC-associated parameters, which are searched for in an unrestricted region of non-negative parameter space. Some biologically unlikely things happen when we look at the QMC parameters - a good example of this are the best-fit values of the viral decay rate *δ*_*V*_ in Fig. 3. For Mouse 2 and 3 (which we see in Fig. 2 are successfully treated by the experimental treatment protocol), QMC predicts that the optimal parameter set has *δ*_*V*_ = 0. Biologically, this at least partially attributes treatment success to the fact that the injected OVs never decay. While it certainly seems reasonable that if the treatment was never eliminated from the body, the tumor volume would go to zero, having no viral decay does not make biological sense. So even though the QMC parameter sets give better fits to the data, the guarantee of a biologically-reasonable value for each parameter is sacrificed.

Looking across methodologies, parameter disparities are the most pronounced in *c*_*T*_, the rate of cytotoxic T cell stimulation from 4-1BBL and IL-12. The QMC-predicted parameters cover a much larger range of values relative to the average mouse. According to the QMC fits, *c*_*T*_ can range anywhere from 92.15% below the value in the average mouse to 4.69 times higher than the value in the average mouse. Compare this to the NLME-predicted values of *c*_*T*_, which can range from 90.29% below the value in the average mouse to 31.87% below the value for the average mouse. What is clear from looking at the best-fit parameter values across methodologies is that it is not differences in a single or small set of parameter values that explain the difference in fits. The nonlinearities in the model simply do not allow the effects of one parameter to be easily teased out from the effects of the other parameters.

### Personalized Treatment Response at Experimental Dose

Here we seek to determine if the two sets of best-fit parameters for a single individual yield similar personalized predictions about tumor response to a range of treatment protocols. The treatment protocols we consider are modeled after the experimental work in [33]. Each day consists of only a single treatment, which can be either an injection of Ad/4-1BBL/IL-12 at 2.5 × 10^9^ viruses per dose, or a dose of 10^6^ DCs. Treatment will be given for six consecutive days, with three days of treatment being Ad/4-1BBL/IL-12, and three days being DCs. If only one dose can be given per day, there are exactly 20 treatment protocols to consider. The 20 protocols are shown on the vertical axis in Fig. 4, where *V* represents a dose of Ad/4-1BBL/IL-12, and *D* represents a dose of dendritic cells.

**Fig 4.**
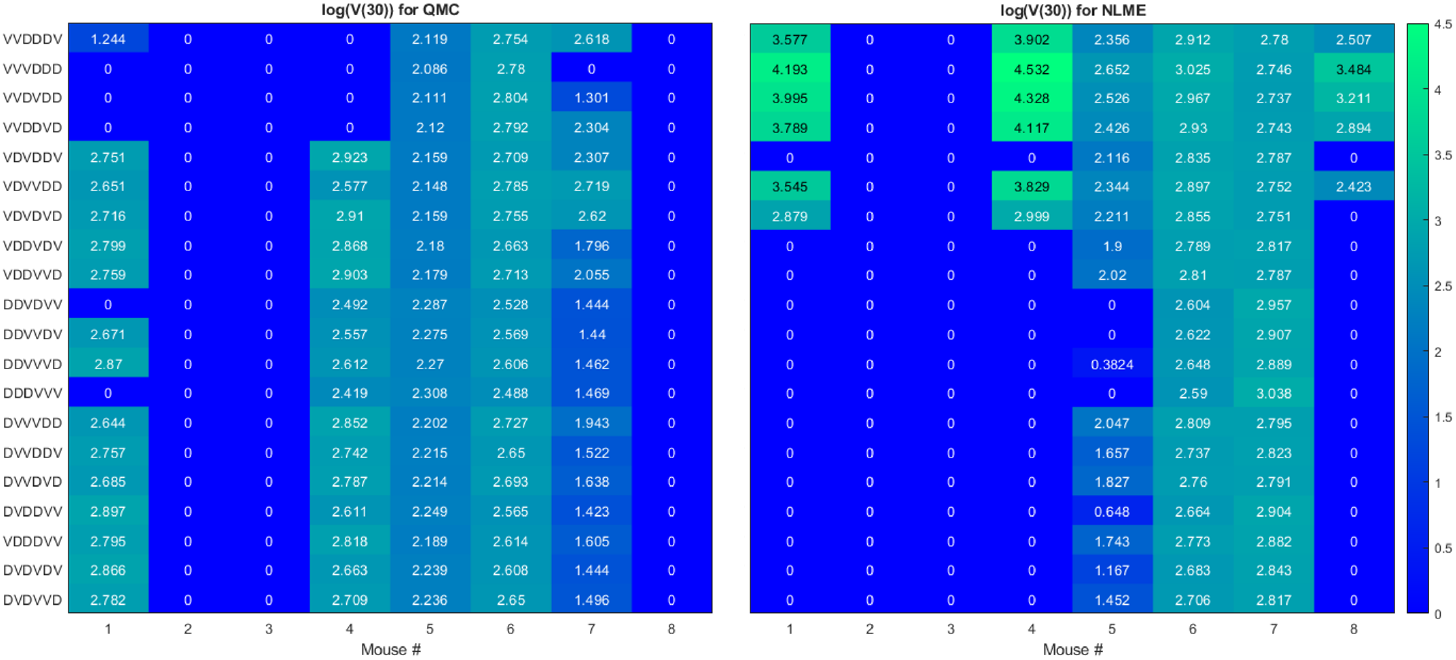
Heatmaps showing the log of the tumor volume measured at 30 days, at the OV and DC dose used in [33]. If log(*V* (30)) ≤ 1, its value is shown as 0 on the heatmap. Left shows predictions when parameters are fit using QMC, and right shows NLME predictions.

To quantify predicted tumor response, we will simulate mouse dynamics using the determined best-fit parameters for each of the 20 6-day protocols. Unless otherwise stated, we will use the predicted tumor volume after 30 days, *V* (30), to quantify treatment response. For each fitting methodology, mouse, and protocol we display the log (*V* (30)) in a heatmap (as in Fig. 4). For all *V* (30) ≤ 1 mm^3^, we display the logarithm as 0, as showing negative values would hinder cross-methodology comparison and overemphasize insignificant differences in treatment response. We consider all such tumors to be effectively treated by the associated protocol. Any nonzero values correspond to the value of log (*V* (30)) when *V* (30) > 1 mm^3^, and we assume these tumors have not been successfully treated. The resulting heatmap at the experimental dose of 2.5 × 10^9^ viruses per dose, and 10^6^ DCs per dose is shown in Fig. 4.

Ideally, we would find that treatment response to a protocol for a given mouse is independent of the fitting methodology utilized, at least in the qualitative sense of treatment success or failure. However, that does not generally appear to be the case for our data, model and fitting methodologies, as we elaborate on here.

- **Cumulative statistics on consistencies across methodologies**. As shown in Fig. 4, the two fitting methodologies give the same qualitative predictions for 73.75% (118/160) of the treatment protocols. Of the 118 agreements, 57 consistently predict treatment success whereas 61 consistently predict treatment failure. It is of note that these numbers only change slightly if we use *V* (80) as our measurement for determining treatment success or failure (81.875% agreement with 78/131 consistently predicting eradication and 53/131 consistently predicting failure - see Fig. S3). Mouse 2, 3 and 6 have perfect agreement across fitting methodologies, and Mouse 7 has 95% agreement across methodologies. For these mice, treatment response is generally not dependent on dosing order. For instance, Mouse 2 and 3 are successfully treated by all twenty protocols considered, whereas Mouse 6 cannot be successfully treated by any protocol. In fact, *V* (30) for Mouse 6 is highly conserved across dosing order, suggesting that the ordering itself is having minimal impact on treatment response. While performing a bifurcation analysis in 11D parameter space is not feasible, what is clear is that for the mice with significant agreement across methodologies, the best-fit parameters must be sufficiently far from the bifurcation surface, as shown in the schematic diagram in Fig. 5. As a result, predicted treatment response is not sensitive to changes in the parameter values that result from using a different fitting methodology. While not equivalent, they also do not appear to be sensitive to dosing order.
- **Cumulative statistics on inconsistencies across methodologies**. The two fitting methodologies give different qualitative predictions for 26.25% (42/160) of the treatment protocols (see Fig. 4). Mouse 1 and 4 are largely responsible for these predictive discrepancies, with Mouse 1 having inconsistent predictions for 75% of protocols, and Mouse 4 having inconsistent predictions for 90% of protocols. Note that each methodology must agree for the protocol VDVDVD, as this was the experimental protocol that was used for parameter fitting. So, 95% is the maximum disagreement rate we can see across methodologies for a given mouse. We observe that the QMC-associated parameter set is much more likely to predict treatment failure for these mice, whereas the NLME parameter set is more likely to predict treatment success. Contrary to the mice for which there is significant cross-methodology agreement, we see a high dependency of treatment response to dosing order for Mouse 1 and 4. From the perspective of the high dimensional bifurcation diagram, these parameters must fall sufficiently close to the bifurcation surface so that parametric changes that result from using different fitting methodologies can lead to wildly different predictions about treatment response (see schematic in Fig. 5). In turn, this appears to make these mice significantly more sensitive to dosing order.

Though the results in this paper are presented for one best-fit parameter set per methodology, we have also explored how parametric uncertainty influences treatment predictions. In particular, for the QMC fitting method, for each mouse we identified suboptimal parameter sets by performing Sobol sampling in a 10% range about the optimal parameter set. Any sampled parameter set that gives a goodness-of-fit within 10% of the optimal is considered a suboptimal parameter set (see Fig. S4). For all such suboptimal parameter sets, treatment response to the 20 protocols was determined. This allows us to study if binary treatment response is insensitive to the precise best-fit parameters used. In Fig. S5 we show the probability a treatment is effective for each mouse across all suboptimal parameter sets. Overwhelmingly, treatment response predicted for an individual mouse and protocol shows excellent agreement across suboptimal parameter sets. Besides treatment response to the protocols VDVVDD and VDVDVD for Mouse 9, predicted treatment response across suboptimal parameter sets agrees over a minimum of 95% of the suboptimal parameter sets. This is seen in Fig. S5 by the probabilities of an effective treatment being either > 0.95 or < 0.05. For this reason, we conclude it is reasonable to compare the prediction across methodologies using only the best-fit parameters.

**Fig 5.**
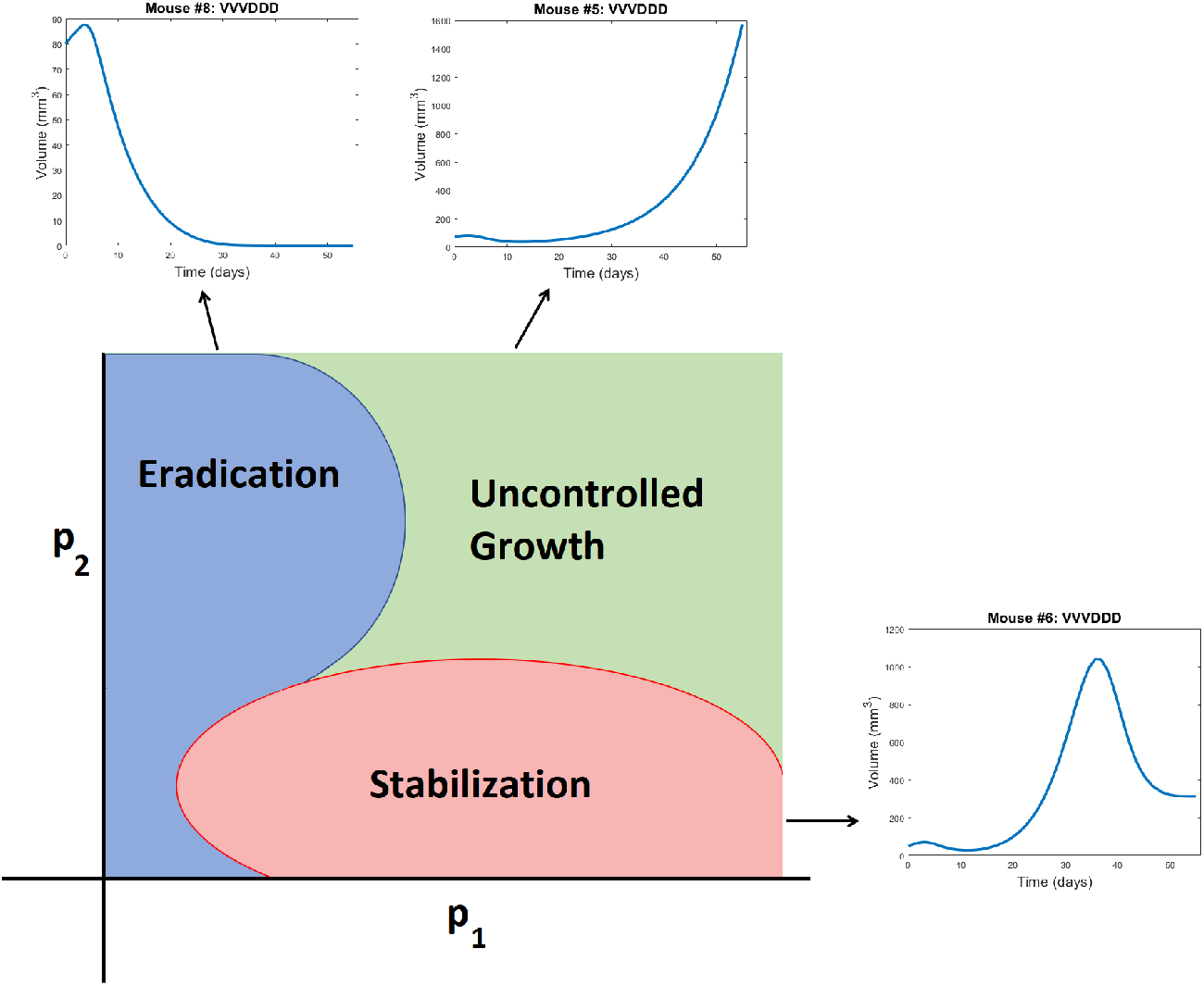
Schematic representation of a bifurcation diagram in two-dimensional parameter space. For certain nonlinear combinations of parameters, a treatment can successfully eradicate a tumor (as occurs for Mouse 8 treated with VVVDDD according to NLME parameters), result in tumor stabilization (as occurs for Mouse 6 treated with VVVDDD according to NLME parameters), or can fail to control the tumor (as occurs for Mouse 5 treated with VVVDDD according to NLME parameters). Note the bifurcation diagram is dependent on both the dose of drug being given, and the ordering of those drugs.

### Exploring Predictive Discrepancies between Fitting Methodologies

The predictive discrepancies across fitting methodologies begs the question of whether the parameters we are fitting are actually practically identifiable given the available experimental data. To explore this question, we generated profile likelihood curves for fitting the *average* tumor growth data. Parameter fitting follows the QMC algorithm presented in the subsection Independently Fitting Individuals, with the exception that the cost function utilizes the average volume 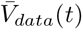 and incorporates the variance in the volume across mice (*σ*^2^(*t*)):

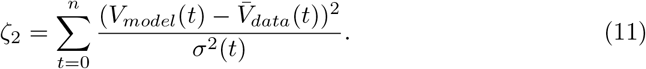

As a first step, we fixed the parameters whose values we could reasonably approximate from experimental data: *δ*_*I*_ = 1, *α* = 3000, *δ*_*V*_ = 2.3, *κ*_0_ = 2, *δ*_*T*_ = 0.35, and *δ*_*D*_ = 0.35 [36]. This means we are using *df* = 5 in the calculation of the threshold, as the generation of each profile likelihood curve requires fitting four model parameters, and the initial condition *U* (0).

The resulting profile likelihood curves in Fig. 6 show that, even under the assumption that six of the eleven non-initial condition parameters are known, several of the fit model parameters lack practical identifiability. The tumor growth rate *r* and the infectivity parameter *β* are both practically identifiable, ignoring slight numerical noise. The T cell activation parameters *χ*_*D*_ and *c*_*T*_ lack practical identifiability as they have profiles with a shallow and one-sided minimum [42]. The profile for *c*_*kill*_ demonstrates that the model can equally well-describe the data over a large range of values for this enhanced cytotoxicity parameter. The flat likelihood profile is indicative of (local) structural unidentifiability, which also results in the parameter being practically unidentifiable [42]. It is worth noting that the original work fitting to the average mouse was done in a *hierarchical* fashion [34, 36], and this circumvented the identifiability issues that emerge when doing simultaneous parameter fitting.

**Fig 6.**
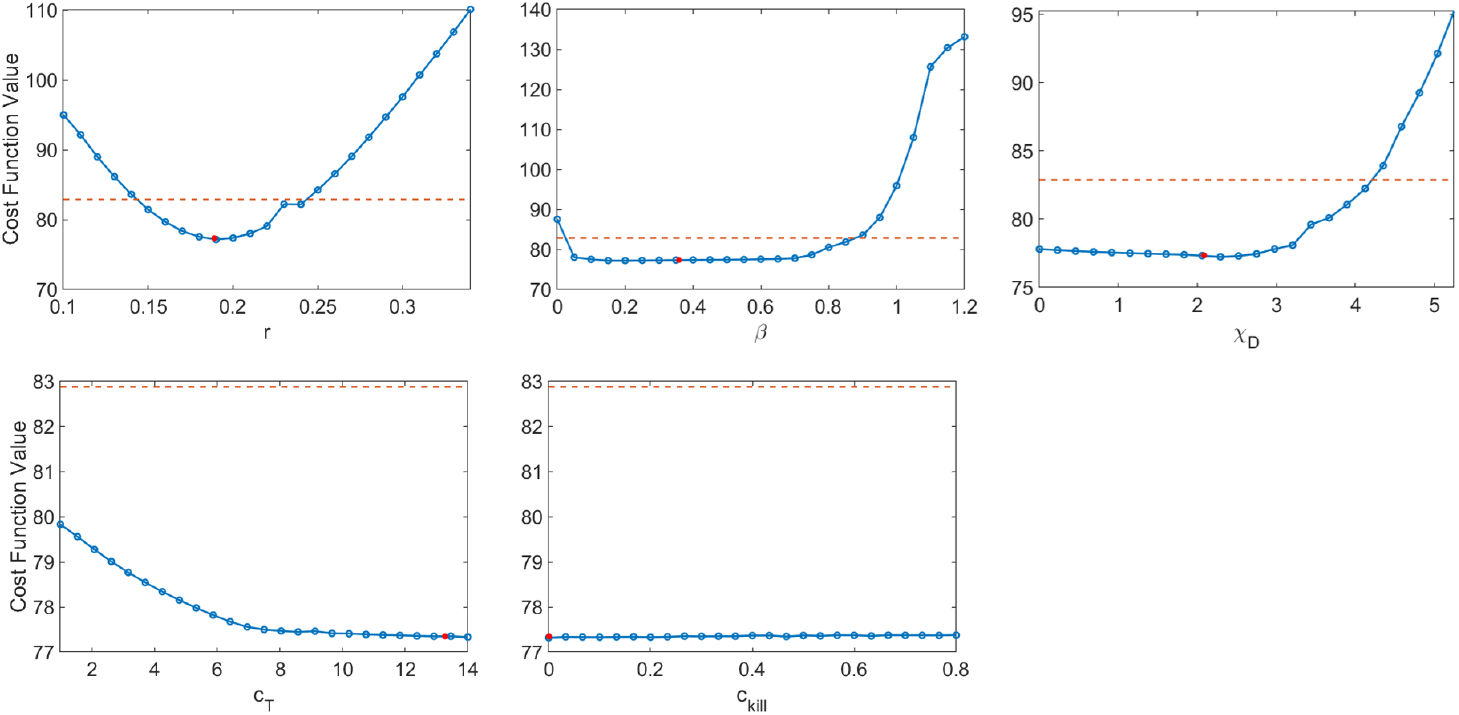
Profile likelihood curves. Top row: tumor growth rate *r*, infectivity rate *β*, T cell activation rate by DCs *χ*_*D*_. Bottom row: T cell stimulation rate by immunostimulants *c*_*T*_, and rate at which immunostimulants enhance cytotoxicity of T cells *c*_*kill*_. The threshold (red dashed line) is calculated using *df* = 5 and a 95% confidence interval.

As we are unable to exploit the benefits of hierarchical fitting when performing personalized fits, this lack of practical identifiability poses significant issues for treatment personalization. We have already seen the consequences of this when we observed that despite both giving good fits to the data, QMC and NLME make consistent qualitative predictions in only 73.75% of the treatment protocols tested across all individuals. While the lack of practical identifiability helps explain why this can happen, it does not explain the mechanisms that drive predictive differences. To this end, we will now focus on the simulated dynamics of Mouse 4 in more detail, as this was the mouse with the most predictive discrepancies across methodologies.

As shown in Fig. 7, when we simulate the model ten days beyond the data-collection window, we see that the QMC and NLME parameters fall on different sides of the bifurcation surface. In particular, in the QMC-associated simulation, at around 34 days the tumor exhibits a local maximum in volume and continues to shrink from there (Fig. 7, left). This is in comparison to the NLME-associated simulation, where the tumor grows exponentially beyond the data-collection window. To uncover the biological mechanism driving these extreme differences, we look at the “hidden” variables in our model - that is, variables for which we have no experimental data. As shown in Fig. 7, despite the similar fits to the volumetric data, the two parameters sets predict drastically different dynamics for the OVs and T cells. For the NLME-associated parameters, the virus and T cell population die out, eventually resulting in unbounded tumor growth. On the other hand, the virus and T cell population remain endemic throughout the simulation when using the QMC-associated parameters, driving the tumor population towards extinction.

**Fig 7.**
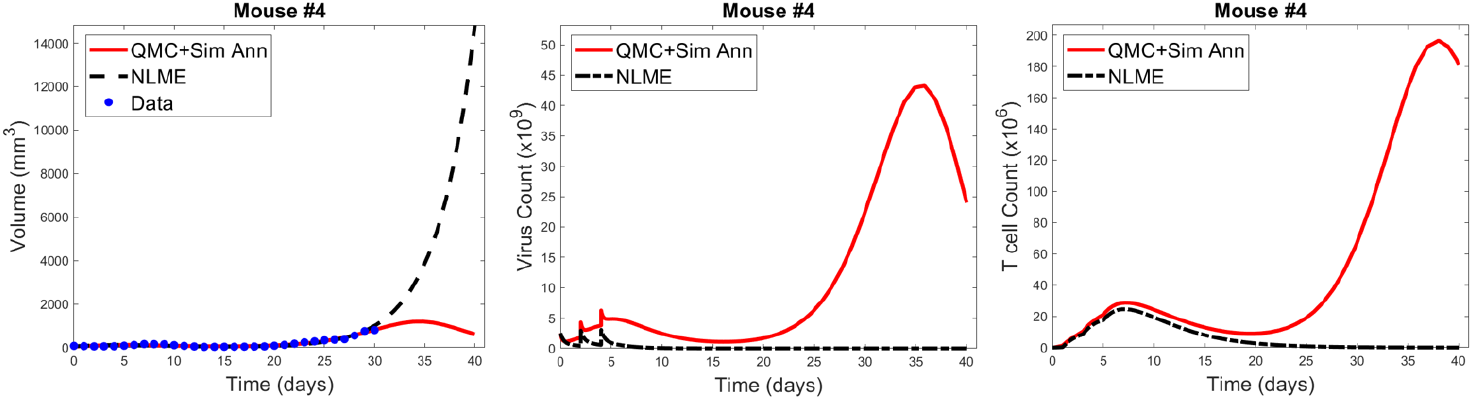
Left: QMC and NLME-associated fits to Mouse 4 treated with VDVDVD, with model predictions extended 10 days beyond the data-collection window. Center and right: Predicted virus and T cell counts associated with each fitting methodology, respectively.

It is common knowledge that more data improves parameter identifiability. Not all data is created equal, however. We could get a lot more time-course data on total tumor volume over the 30-day window, but that would not necessarily improve parameter identifiability. Instead, we have identified that the addition of a single data point, for the right variable, at the right time, could go a long way in reconciling predictive discrepancies across fitting methodologies. To make this concrete, suppose we had data that, for Mouse 4, no tumor-targeting T cells are detected at 30 days. If we introduced a modified cost function that weighed both the contribution of the tumor volume and this T cell measurement, the parameter set identified by QMC would no longer be optimal, as it predicts a T cell burden on the order of 10^8^ (100 × 10^6^). The optimal parameter set should be one for which *T* → 0, and once this occurs, there is no mechanism to control the tumor in the long-term. As a result, the tumor must regrow, just as predicted for the NLME-associated parameters. While this thought experiment does not suggest all practical identifiability issues would be reconciled by having this one data point, it does indicate why the predictive discrepancies we see for Mouse 4 (and also Mouse 1) would be at least partially resolved by the addition of a single data point on tumor-targeting T cell volume. This highlights that although one must be quite cautious in using mathematical models to make personalized predictions, models can help us determine precisely what additional data is needed so that we can have more trust in our mathematical predictions.

### Personalized Treatment Response to the Optimal for Average Protocol

Ideally, when an optimal prediction is made for the average of a population, that optimal treatment protocol would also well-control the tumors of individual patients in the population. However, it is well known and supported by our earlier work with virtual populations that this is not necessarily the case. In [31] we showed that the experimental dose being considered in this paper is *fragile or non-robust*. We define a dosing regime as fragile if virtual populations that deviate somewhat from the average population have the same qualitative response to the optimal-for-the-average protocol. In particular, while VVVDDD was the optimal-for-the-average of the mice in the experiments (and this optimal led to tumor eradication for the average mouse [36]), we found only 30% of virtual populations were successfully eradicated by this protocol [31]. Importantly, fragility is a probabilistic population-level descriptor, and not an individual descriptor. While it tells us that populations that deviate somewhat from the average are less likely to behave like the average, it tells us nothing about individuals, particularly if the individuals have behavior that deviates significantly from the average (which is often the case, as shown in Fig. 1). Though, it seems natural to hypothesize that in such a fragile region we may have to be more careful about applying a prediction for the average of a population to any one individual in that population.

We will explore that hypothesis here by looking at statistics on how individual mice respond to VVVDDD, the predicted optimal treatment protocol for the average mouse. While this protocol was effectively able to eradicate the average tumor in the population, its success across individual mice varies significantly across fitting methodologies. For the QMC-associated predictions, this protocol eradicates tumors in 75% of the individual mice (second row of the heatmaps in Fig. 4, left). Compare this to the NLME-associated predictions, in which this protocol eradicates tumors in only 25% of the individual mice (second row of the heatmaps in Fig. 4, right). As shown in Fig. S3, this prediction is unchanged if we determine treatment success or failure at day 80 instead of day 30.

We can also compare response to the optimal-for-the-average protocol across methodologies. We see a qualitative agreement across methodologies (eradication or treatment failure) in only 50% of the mice (Mouse 2, 3, 5, 6). Mouse 7 is particularly interesting, as there was 95% agreement across methodologies when using *V* (30) to measure treatment success or failure, and the optimal for the average of VVVDDD is the only protocol for which treatment response differed (with QMC predicting tumor eradication, and NLME predicting treatment failure). As a further sign of caution, notice how for Mouse 1 and 4 (the cases with significant predictive discrepancies across methodologies), and Mouse 8 (intermediate case with 25% predictive discrepancies), VVVDDD eradicates the tumor with the QMC-associated parameters yet is the **worst protocol** that could be given (largest log (*V* (30))) for the NLME-associated parameters. This is particularly unsettling as it means the population-level optimal treatment recommendation could be the worst-case scenario for some individuals. This confirms our hypothesis that a population-level prediction should be applied to individuals very cautiously when in a fragile region of dosing space.

This raises the question: what if we were assessing individualized response to the average protocol in a robust region of dosing space? We define a dosing regime as robust if virtual populations that deviate somewhat from the average probabilistically have the same qualitative response to the optimal-for-the-average protocol. In [31] we showed that the high DC (50% greater than experimental dose), low OV (50% lower than experimental dose) region of dosing space is robust, with 84% of virtual populations being successfully eradicated by the optimal-for-the-average protocol of DDDVVV. This probabilistic population-level assessment of robustness naturally leads to the hypothesis that in a robust region of dosing space, we may have more success with the optimal-for-the-average treatment in individual mice. We will explore that hypothesis here.

The robust population-level optimal of DDDVVV yields a successful treatment response in all eight mice for the NLME-associated parameters. This holds whether we use *V* (30), our original measure for establishing treatment success (as shown in Fig. S6), or if we use *V* (80) as shown in Fig. 8. This is consistent with the robust nature of this region of dosing space, as the NLME-associated parameters are less likely to wildly deviate from the average mouse due to population-level distributions constraining the value of these parameters. In comparison, the QMC-associated predictions show that only 62.5% of the individual mice are successfully treated by the optimal for the average in an 80-day window (Fig. 8, top left). That said, if we look at the data more closely, we can see that Mouse 7 has essentially been eradicated even though 80 days was not quite long enough to drive *V* (80) < 1 mm^3^, our threshold for eradication. Fig. 8 also shows that the tumor volume for Mouse 6 has stabilized. Thus, we see that the QMC-associated predictions actually agree with the optimal-for-the-average response in 75% of cases (or, 87.5% if you consider the stabilization of Mouse 6 to be a “success” rather than a “failure”).

**Fig 8.**
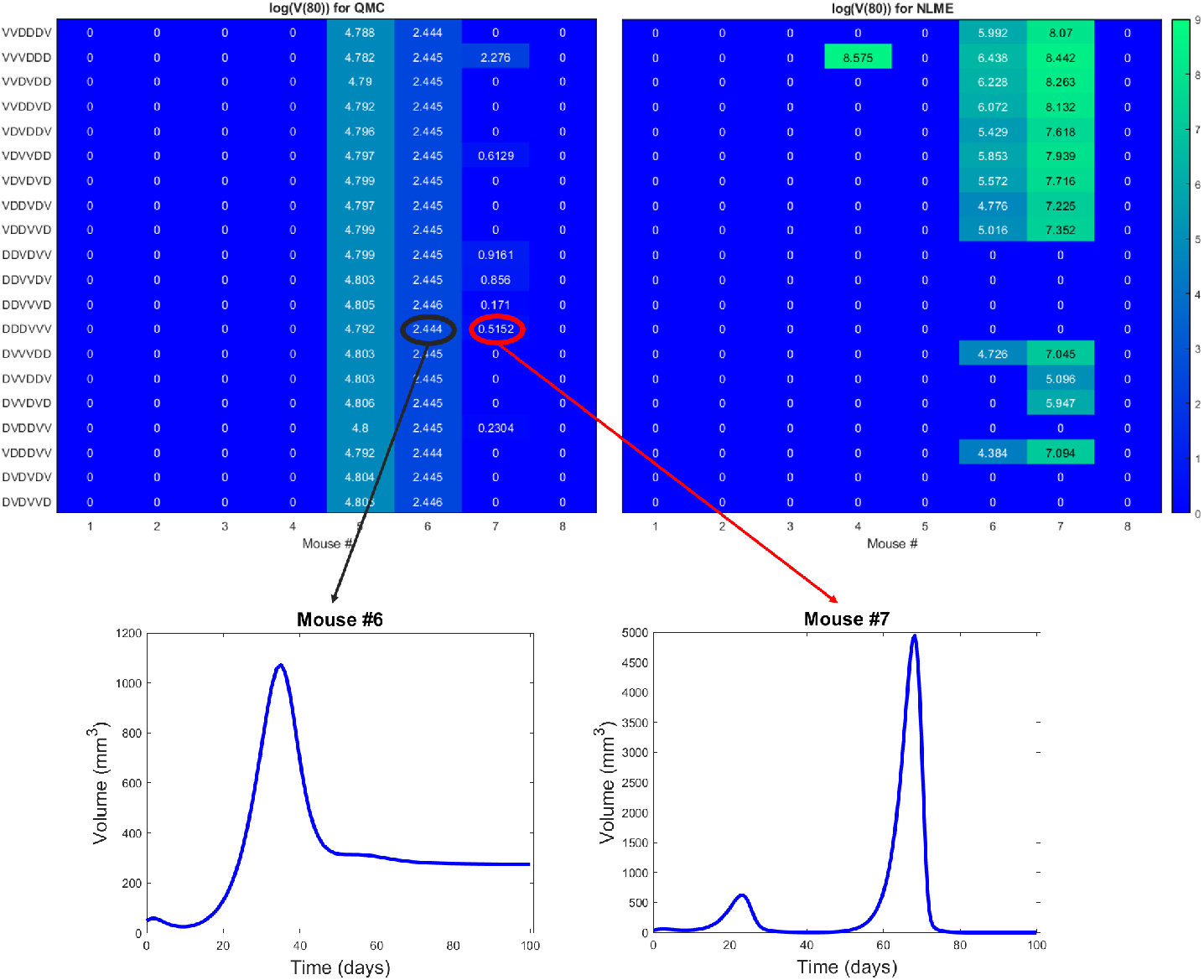
Heatmaps showing the log of the tumor volume measured at 80 days, at the high DC (50% greater than experimental dose), low OV (50% lower than experimental dose) region of dosing space. Left shows predictions if parameters are fit using QMC, and right shows NLME predictions. Inserts show time course of predicted treatment response for Mouse 6 and 7 to the optimal-for-the-average protocol of DDDVVV.

In closing, we have confirmation of our hypothesis that there is a significant benefit to working with a robust optimal-for-the-average protocol, even in the absence of all model parameters being practically identifiable. In the presence of robustness, we predict that one could generally apply the optimal-for-the-average protocol and expect a qualitatively similar response in most individuals. While this does not mean each individual is treated with their personalized optimal protocol, this has important consequences for determining when a population-level prediction will be effective in an individual.

## Conclusion

In this work, we demonstrated that computational challenges can arise when using individualized model fits to make treatment recommendations. In particular, we showed that treatment response can be sensitive to the fitting methodology utilized when lacking sufficient patient-specific data. We found that for our model and preclinical dataset, predictive discrepancies can be at least somewhat explained by the lack of practical identifiability of model parameters. This can result in the dangerous scenario where an effective treatment recommendation according to one fitting methodology is predicted to be the worst treatment option according to a different fitting methodology. This raises obvious concerns regarding the utility of mathematical models in personalized oncology.

While it is well-established that more data improves parameter identifiability, here we highlight how we can identify precisely what data would improve the reliability of model predictions. In particular, we see how having a single additional measurement on the viral load or T cell count at the end of the data collection window would go a long way to reduce the predictive discrepancies across fitting methodologies (Fig. 7). When additional data is not available, an alternative option to personalization is simply treating with the population-level optimal. Here we showed the dangers of applying the optimal-for-the-average for a fragile protocol, and we demonstrated that such a one-size-fits all approach is much safer to employ for a robust optimal protocol. Therefore, even when data is lacking to make personalized predictions, establishing the robustness of treatment response can be a powerful tool in predictive oncology.

It is of note that this study uses just one mathematical model, with one set of assumptions, to reach our cautionary conclusion regarding the fitting methodology utilized and the resulting biological predictions. And this model is quite a simple one, ignoring many aspects of the immune system, and spatial aspects of immune infiltration (as done in [48], among many other references). The model used herein was chosen because it has been previously validated against the average of the available experimental data. A more complex model would be problematic here, as there is simply not the associated experimental data to validate such a model. While this study certainly does not guarantee that similar issues will arise when working with other models and datasets, it highlights the need for caution when using personalized fits to draw meaningful biological conclusions.

As we enter the era of healthcare where personalized medicine becomes a more common approach to treating cancer patients, harnessing the power of mathematical models will only become more essential. Understanding the identifiability of model parameters, what data is needed to achieve identifiability, and whether treatment response is robust or fragile are all important considerations that can greatly improve the reliability of personalized predictions made from mathematical models.

## Acknowledgments

J.L.G would like to thank Dr. Joanna Wares and Dr. Eduardo Sontag for the many discussions that helped to develop the ideas in this manuscript. The authors are also grateful to Dr. Chae-Ok Yun for the exposure and access she provided to the rich dataset utilized in this work. E.N. would like to acknowledge Dr. Yang Kuang for his support at the early stages of this project. J.L.G and M.C.L. acknowledge use of the ELSA high-performance computing cluster at The College of New Jersey for conducting the research reported in this paper.

## Supporting information

**Fig S1.**
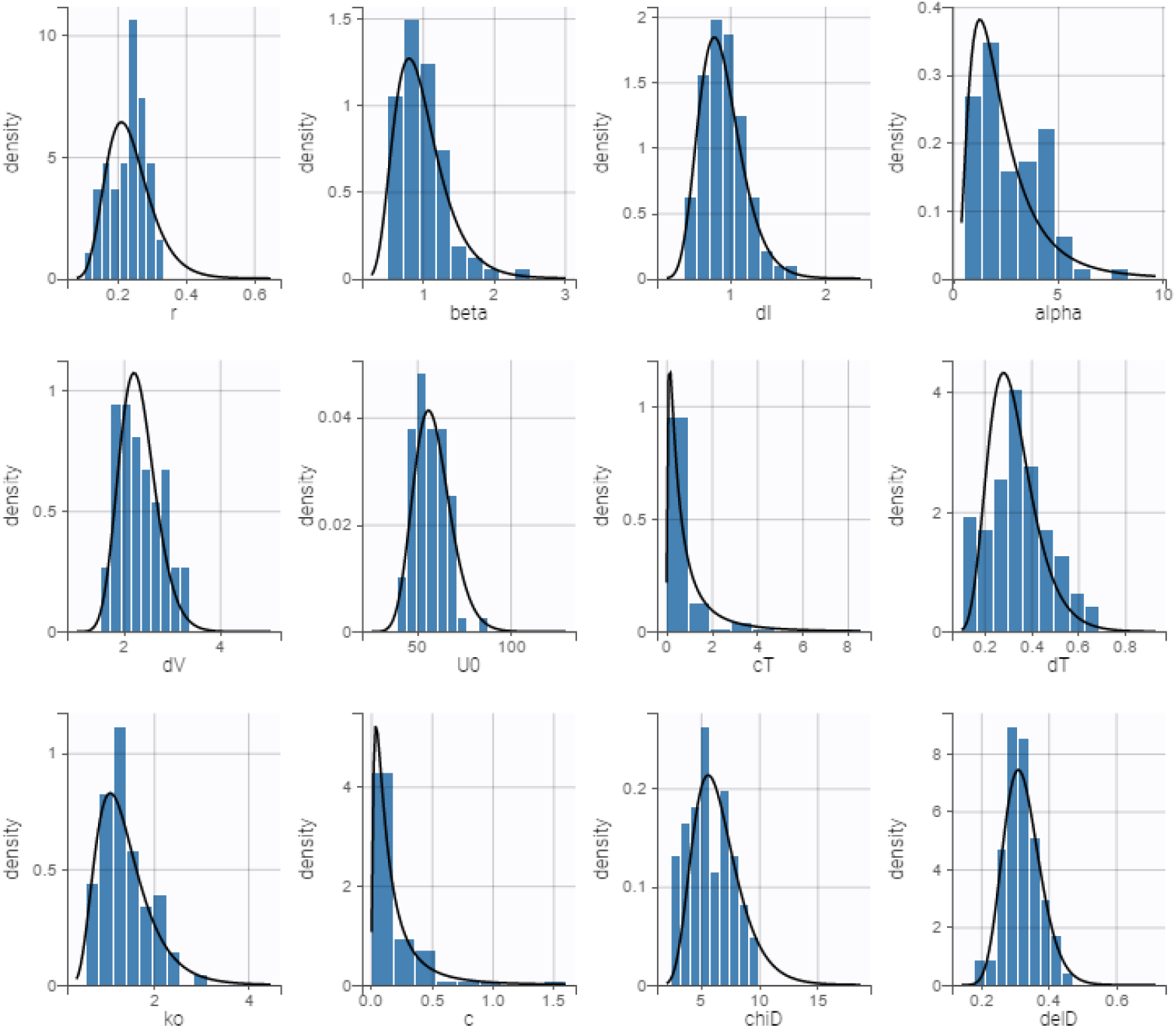
Estimated parameter distributions from Monolix’s implementation of NLME. Each parameter is assumed to be lognormally distributed. The blue bars in each graph represent the empirical distribution of the parameter estimation and the black line represents the theoretical distribution.

**Fig S2.**
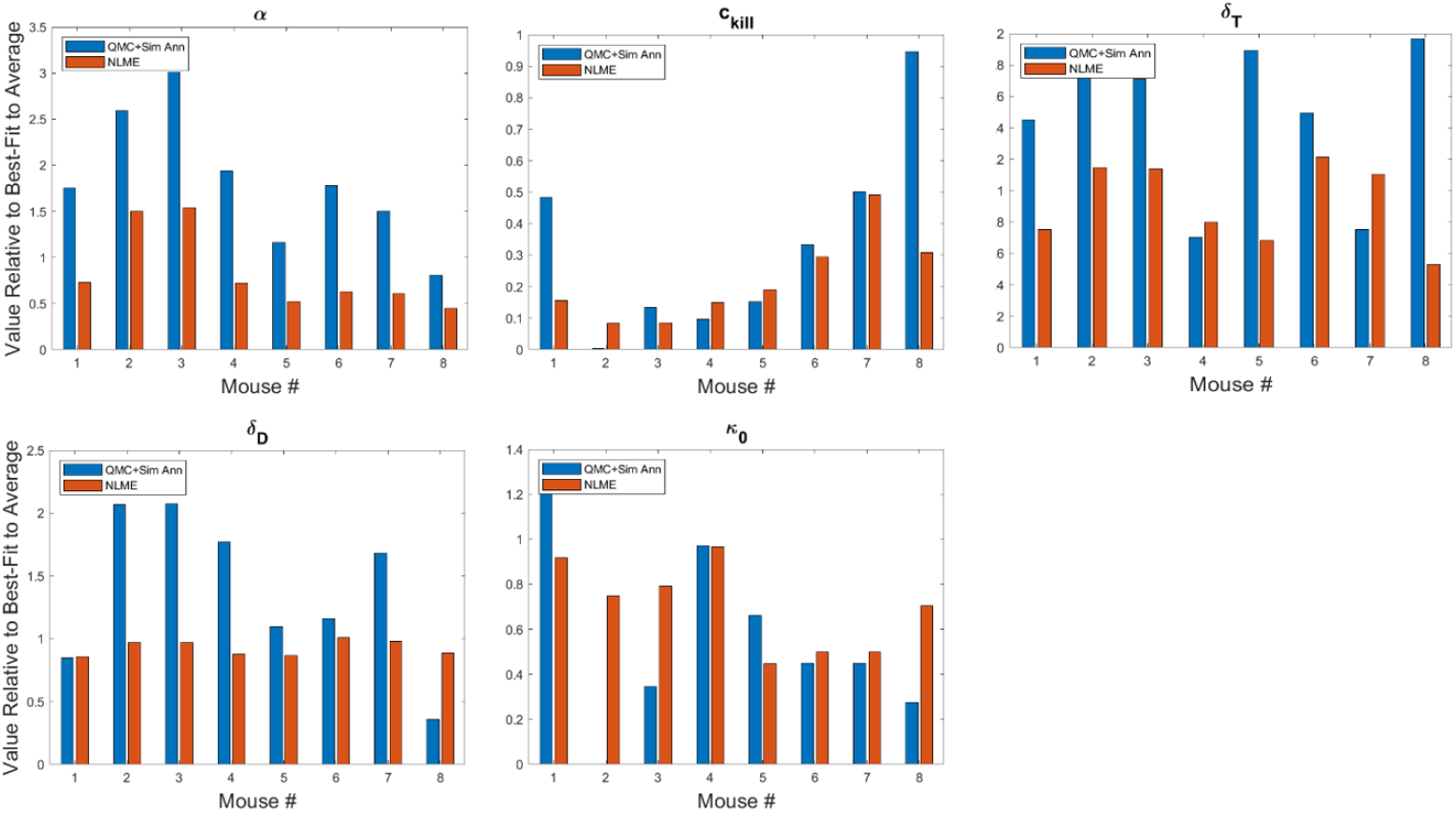
Best-fit value of number of viruses released by lysed cell *α*, the cytotoxicity enhancement term due to immunostimulants *c*_*kill*_, T cell decay rate *δ*_*T*_, DC decay rate *δ*_*D*_, and default cytotoxicity rate of T cells *κ*_0_. The best-fit values are shown for each mouse and are presented relative to the best-fit value of the parameter in the average mouse [34]. Therefore, a value of 1 means the parameter value is equal to that in the average mouse, less than 1 is a smaller value, and greater than 1 is a larger value.

**Fig S3.**
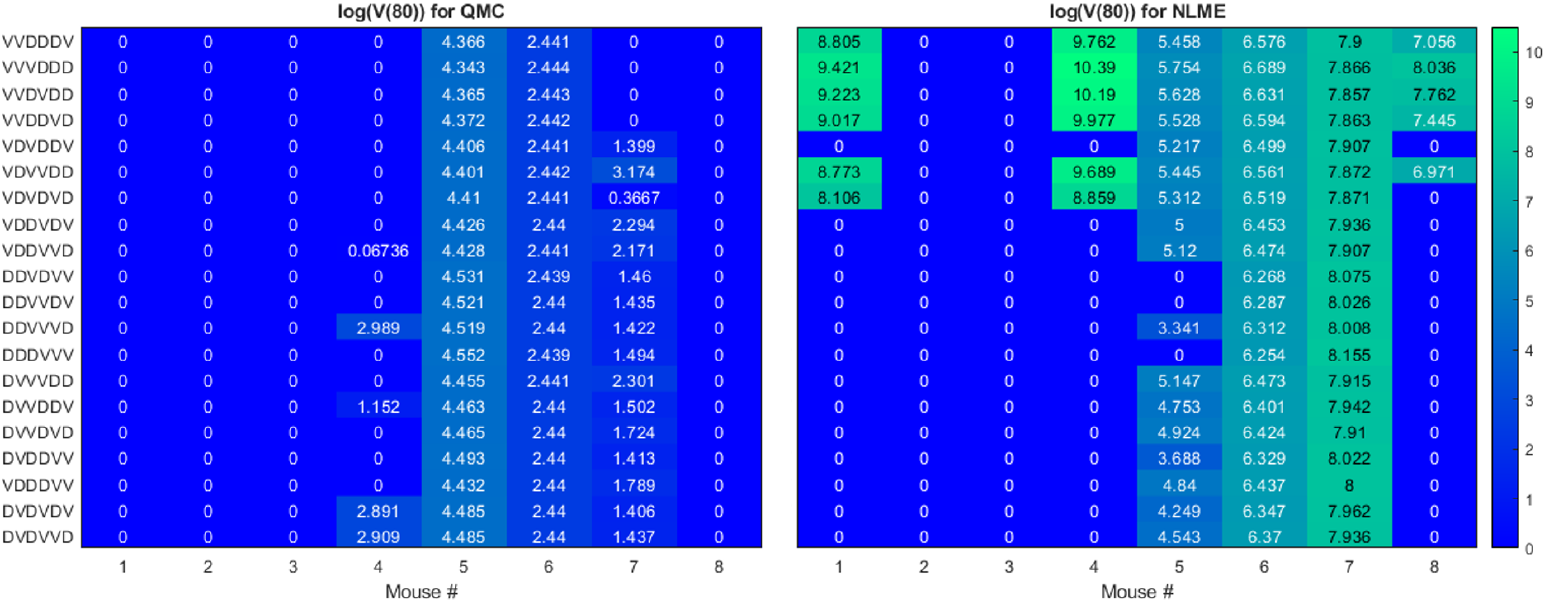
Heatmaps showing the log of the tumor volume measured at 80 days, at the OV and DC dose used in [33]. Left shows predictions when parameters are fit using QMC and right shows NLME predictions. Compare to heatmap in Fig. 4 which shows the log of the tumor volume 50 days earlier.

**Fig S4.**
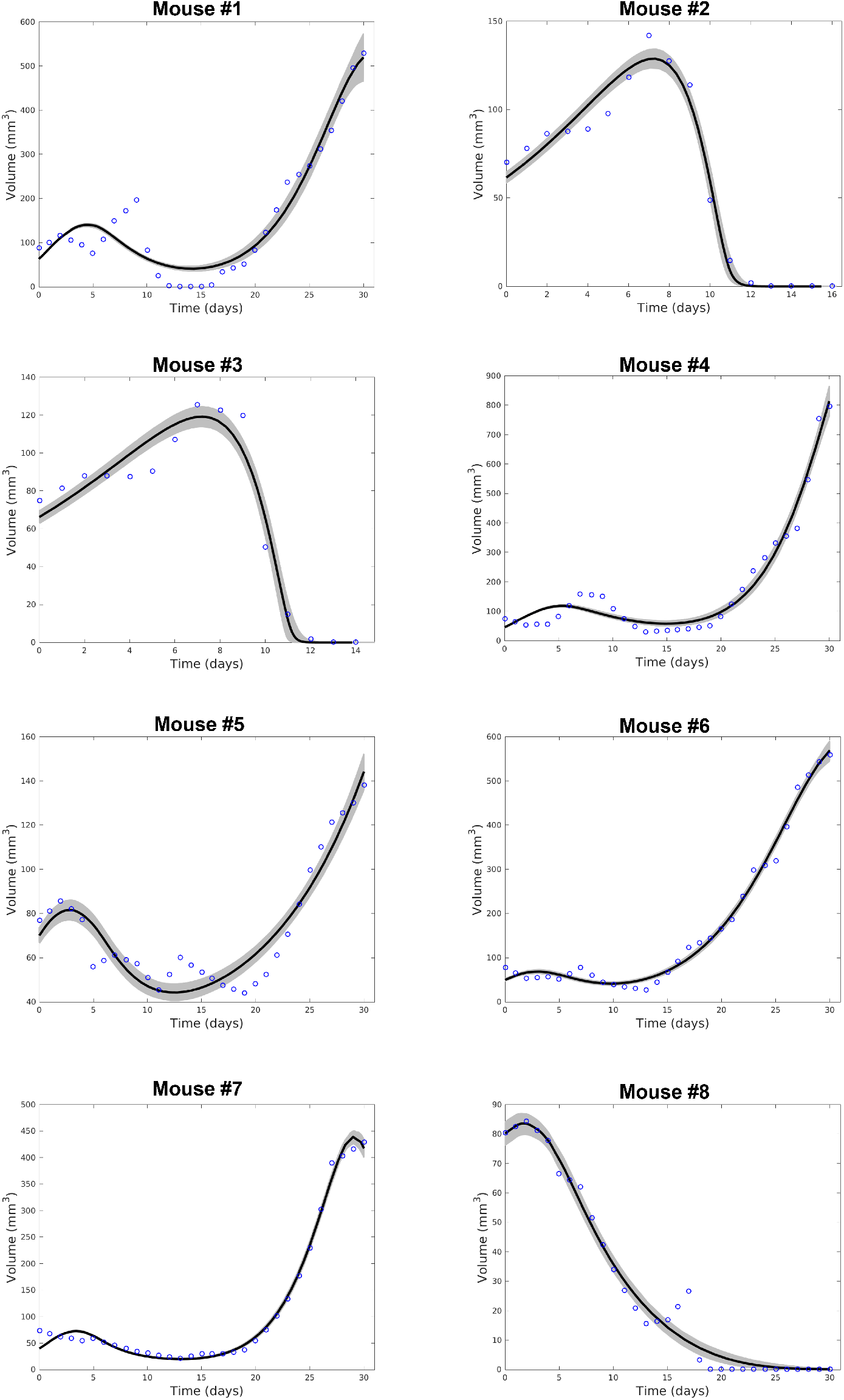
Optimal QMC fit (black) with suboptimal QMC fits (grey) to experimental data (blue outlined circles) for which the goodness-of-fit metric is within 10% of optimal.

**Fig S5.**
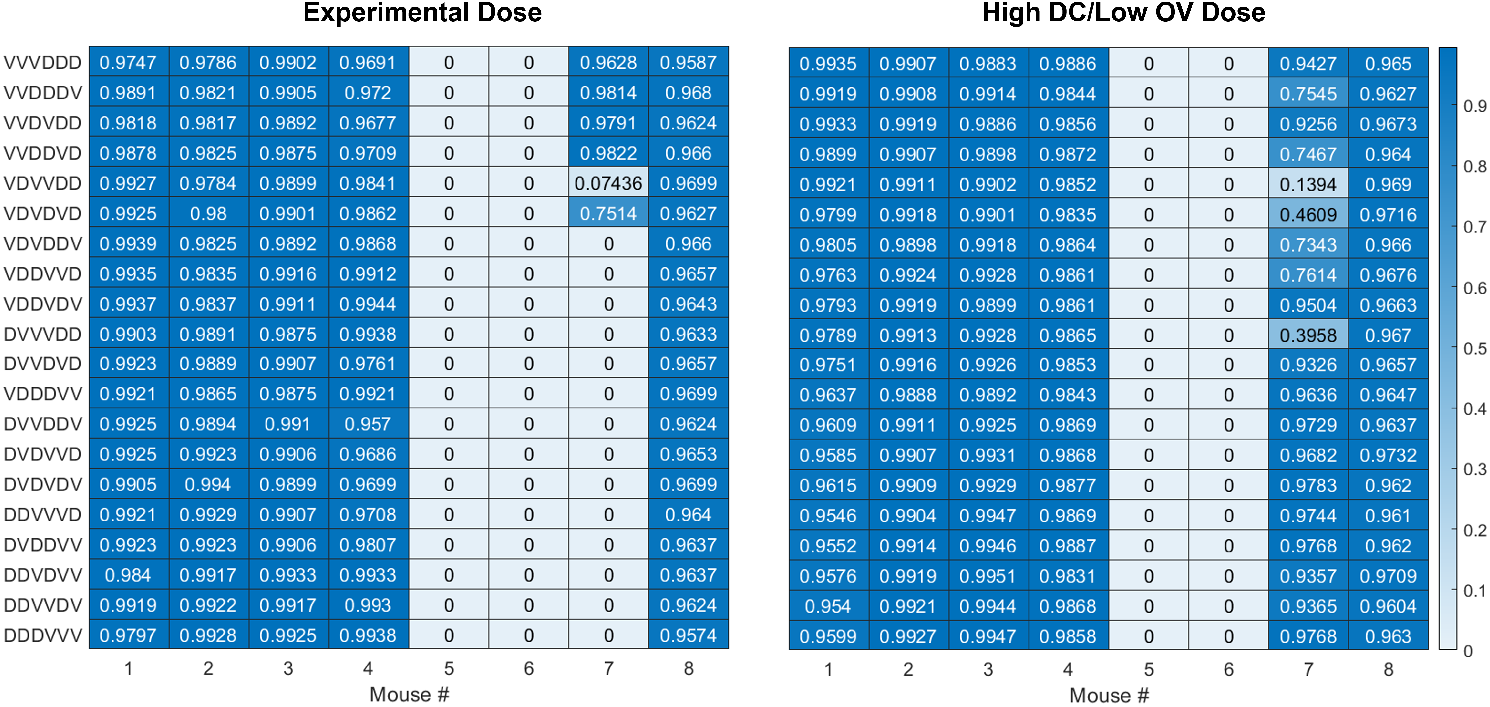
Probability of an effective treatment over the suboptimal parameter sets per mouse, across the 20 treatment protocols. The experimental dose is shown on the left, and the high DC/low OV dose is shown on the right.

**Fig S6.**
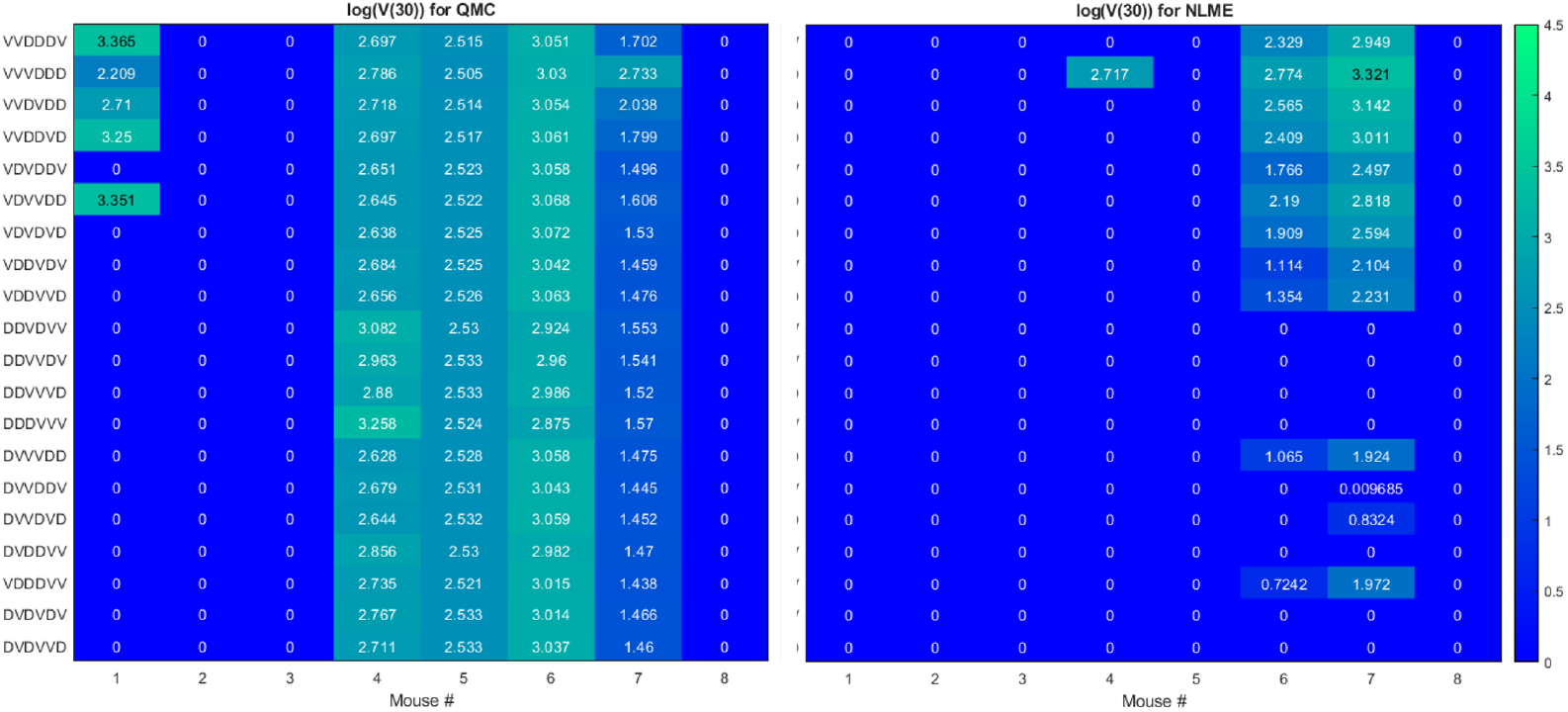
Heatmaps showing the log of the tumor volume measured at 30 days, at the high DC (50% greater than experimental dose), low OV (50% lower than experimental dose) region of dosing space. Left shows predictions if parameters are fit using QMC and right shows NLME predictions. Compare to heatmap in Fig. 8 which shows the log of the tumor volume 50 days later.

## Notes

### Competing Interest Statement

The authors have declared no competing interest.

### Summary of Updates

Changes to title, abstract, introduction and conclusion. Additional results have been added.

